# The effect of venipuncture site on hematology of bats: implications for comparative analyses

**DOI:** 10.1101/2025.02.28.640807

**Authors:** Alicia Roistacher, Bret Demory, Daniel J. Becker

**Author notes:** From the symposium Applied Ecoimmunology in Wildlife Health and Conservation presented at the annual meeting of the Society for Integrative and Comparative Biology, January 3-7, 2025 at Atlanta, Georgia.

## Abstract

Wildlife health comparisons within and across populations and species are essential for population assessment and surveillance of emerging infectious diseases. Due to low costs and high informational yield, hematology is commonly used in the fields of ecoimmunology and disease ecology, yet consistency and proper reporting of methods within the field are lacking. Previous investigations on various wildlife taxa have revealed noteworthy impacts of the vein used for blood collection on hematology measures. However, the impacts of venipuncture site on bats, a taxon of increasing interest in ecoimmunology and disease ecology, have not yet been tested. Here, we use a long-term study system in western Oklahoma to test the effect of venipuncture site on hematology parameters of the Mexican free-tailed bat (*Tadarida brasiliensis*) and cave myotis (*Myotis velifer*), two abundant and representative bat species from the families Molossidae and Vespertilionidae. Between September 2023 and October 2024, we collected paired peripheral blood from both the propatagial and intrafemoral veins in 25 individuals per species. We then measured total red and white blood cells, reticulocyte counts, and leukocyte differentials and used generalized linear mixed models to compare parameters among venipuncture sites within and between bat species. Overall, venipuncture site had no effect on any hematology parameters; however, we revealed small differences in neutrophil and lymphocyte proportions between veins among the species. By contrast, we detected significant species-level differences in most cell measurements, which we propose could be explained by life-history strategy and phylogenetic differences. We encourage continued testing of additional venipuncture sites, and of the same venipuncture sites on different species, on hematology and other health metrics used in ecoimmunology and disease ecology. Lastly, we emphasize the significance of thorough method reporting in publications to enable transparent comparisons and accounting for even small sampling-based artifacts. All future efforts are especially important for bats to improve conservation monitoring, ecosystem services estimations, and their association with emerging infectious diseases.

## Introduction

Hematology is widely used in the fields of ecoimmunology and disease ecology owing to its use of non-lethal, low-cost, and small volume blood samples (Kloskowski et al., 2017; Maceda-Veiga et al., 2015). Peripheral blood from vertebrates reflects whole-organism function and can be subject to many assays, including but not limited to differential white blood cell (WBC) and red blood cell (RBC) counts, blood chemistry panels, platelet counts, comet assays, and pathogen screening (Dziki-Michalska et al., 2024; Gupta & Nigar, 2020; Maceda-Veiga et al., 2015). The data obtained from these assays can provide insights into innate and adaptive immune function, intrinsic and extrinsic stressors, and pathogen infections (Gupta & Nigar, 2020; Kloskowski et al., 2017). Blood smears are the most widely used sample when studying wildlife health due to their low costs and blood requirement relative to their high information yield (Maceda-Veiga et al., 2015). For example, ratios of neutrophils to lymphocytes (NL ratios) are commonly used to assess stress and immunity (Davis et al., 2008; DeAnglis et al., 2024; Tovstukha et al., 2024), while RBCs, reticulocytes (RETs), and WBCs are often used to diagnose disease (Pouletty, 2010) as well as detect blood pathogens (Alharbi et al., 2022; Gupta & Nigar, 2020; Schinnerl et al., 2011), inflammation, and tissue damage (Maceda-Veiga et al., 2015).

The vein used to collect peripheral blood (i.e., venipuncture site) has been shown to have a significant effect on hematological and biochemical parameters in diverse wildlife taxa, including but not limited to turtles (Rodríguez-Almonacid et al., 2022), tortoises (Eshar et al., 2016; Gottdenker & Jacobson, 1995; López-Olvera et al., 2003; Neiffer et al., 2021), sharks (Mylniczenko et al., 2006), deer (Comazzi et al., 2023), and birds (Kern & De Graw, 1978; Sheldon et al., 2008). For example, in the Colombian slider (*Trachemys callirostris*), hematocrit, haemoglobin, and RBC count significantly varied between three vein types (Rodríguez-Almonacid et al., 2022). Meanwhile, other studies have hypothesized that detectable differences in hematology results could be explained by differing venipuncture sites in turtles (Crooks et al., 2023; Gregory et al., 2022). Owing to different protocols and logistical constraints, sample collection and storage methods are not standardized across and within research groups or across seasons, populations, and/or species within groups (Maceda-Veiga et al., 2015; Rodríguez-Almonacid et al., 2022). Testing for the potential effects of venipuncture site on hematology across wildlife taxa more generally is important to account for such sampling artifacts in existing and future analyses (Neiffer et al., 2021). Improving upon method standardization, increased efforts in baseline data collection, and reporting of venipuncture type used will enhance the fields of ecoimmunology and disease ecology and improve inference of comparative analyses.

Bats are increasingly studied in ecoimmunology and disease ecology due to their ecological diversity and association with emerging infectious diseases, especially as some species seem to tolerate virulent viruses with little-to-no pathology (Irving et al. 2021; Becker et al. 2025). Hematological methods have been commonly used to assay the bat cellular immune system and response to both intrinsic and extrinsic factors, given the lack of bat-specific immunological reagents, remote field sites and limited cold chain capacity, and the small blood volumes that can be safely collected from many bats (Ruoss et al., 2019; Schneeberger et al., 2013). Cellular immune profiles significantly vary across bat species due to their diverse ecological characteristics, evolutionary histories, and geographical distributions (Schneeberger et al. 2013; Cornelius Ruhs et al. 2021; Becker et al. 2025). For example, (DeAnglis et al., 2024) revealed annual trends in total and differential WBC counts that varied by bat species within the Neotropics. However, despite increasing research efforts on bat cellular immunity, hematology characterization across the bat phylogeny remains limited (Cornelius Ruhs et al., 2021).

The bat research community has closely followed the universal rule for blood collection across vertebrates: do not collect more than 1% of the animal’s body weight in blood within a 24-hour period (Sikes, 2016; Wolfensohn & Lloyd, 2013). Due to personal preference as well as bat body mass and morphology, different veins are typically used to collect peripheral blood (**Figure 1**). Conventionally, bats are non-lethally bled from either the propatagial (cephalic) or intrafemoral (saphenous) vein, but they are also lethally bled from cranial vena cava, heart, jugular, and orbital sinus veins (Eshar & Weinberg, 2010). This between- and even within-study variation could introduce further noise and biases for comparative analyses and determination of baseline data for newly discovered or underrepresented species. Further, to our knowledge, the effect of venipuncture site on any health metric in bats has not been investigated.

**Figure 1.**
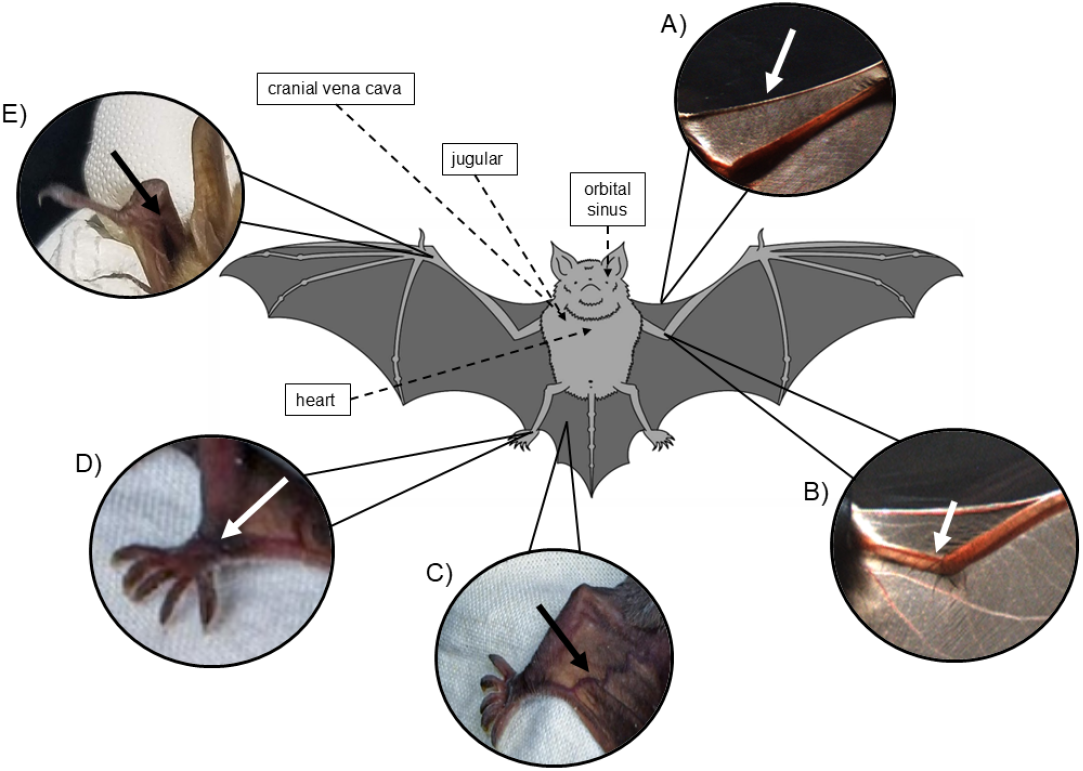
Depiction of all venipuncture sites in bats. The externally visible veins are (A) propatagial (photo: Ralph Simon), (B) brachial (photo: Ralph Simon), (C) intrafemoral (photo: Jon Alonzo), (D) lower intrafemoral (photo: Jon Alonzo), and (E) upper brachial. The other veins that require lethal blood collection are shown with dashed lines.

In this study, we test the effect of the two most commonly used veins for bat blood collection, the propatagial and intrafemoral venipuncture sites, on hematology measures of two sympatric bat species, the Mexican free-tailed bat (*Tadarida brasiliensis*) and cave myotis (*Myotis velifer*). To our knowledge, this represents the first assessment of paired hematology values collected from two venipuncture sites in any bat species. We also contribute baseline hematology measures of these two bat species during critical life-history events for ongoing intra- and inter-specific studies.

## Methods

### Study Species and Sites

We focused our study on two seasonally sympatric bat species in western Oklahoma, as part of a long-term study of bat immunity and infection (Becker et al. 2025). The Mexican free-tailed bat (*Tadarida brasiliensis*), a member of the Molossidae family, migrates from their wintering grounds in central and southern Mexico to the southwestern United States and northern Mexico to give birth and raise their offspring (Villa & Lendell Cockrum, 1962; Wiederholt et al., 2015). In addition to long-distance migration, *T. brasiliensis* form the largest congregations of any mammal, containing up to tens of millions of individuals in per colony (Ammerman et al., 2012; Danielson et al., 2022; Iskali & Zhang, 2015; McCracken et al., 2018). By contrast, the cave myotis (*Myotis velifer*) is part of the Verspertilionidae family and inhabits arid temperate and semi-tropical regions between central Oklahoma to southern California in the United States and into southern Mexico (Krutzsch, 2009). *M. velifer* enter torpor and hibernate during winter (Fitch et al., 1981; Krutzsch, 2009), although they have been observed undergoing short-distance or altitudinal migrations to their hibernacula (Ayala-Berdon & Solis-Cardenas, 2017; Donald W. Tinkle, 1965). Compared to *T. brasiliensis, M. velifer* form smaller colonies of up to 10,000 individuals per group (Humphrey & Oli, 2015), but these are large colonies for the genus *Myotis* (Burrell & Bergeson, 2022; Cheng et al., 2021; Santana et al., 2011). Both species are insectivores, contributing to significant reduction of annual agricultural costs of pesticides (Danielson et al., 2022; Wiederholt et al., 2015). The two species are often found co-roosting in caves and anthropogenic dwellings (Betke et al., 2024) share similar reproductive characteristics, and both form maternity colonies during summer months (Fitch et al., 1981; Wilkins, 1989).

Bats were captured at the Selman Bat Cave and Alabaster Caverns State Park in Woodward County, Oklahoma. Woodward County sits within the Cimarron Gypsum Hills karst area of Oklahoma (William & Samanie, 2010). Both caves have exposed gypsum, are overlain with shale layers, surrounded by sage bush, cattle pasture, and high rolling plains separated by eroded canyons (Bryant, 2013; Caire et al., 1984; William & Samanie, 2010). These sites were *chosen* based on the large populations of *T. brasiliensis* and *M. velifer* inhabiting the caves. The Selman Bat Cave houses an estimated 50,000 *T. brasiliensis* each year over their summer occupancy (Betke et al., 2008), and the cave system we used at Alabaster Caverns State Park houses multiple bat species, including both *T. brasiliensis* and *M. velifer* from spring into early autumn (Caire et al., 1984). Caves in Woodward County have been reported as hibernacula for *M. velifer* (Caire & Loucks, 2010; Humphrey & Oli, 2015). *There is a lack of migration reports for this species in northwest Oklahoma; thus we assume M. velifer* strictly hibernates here.

We sampled bats between September 2023 and October 2024. Bats were captured with hand nets at the Selman Bat Cave and with hand nets and mist-nets at the target cave at the Alabaster Caverns State Park. Both sexes (*M. velifer*: 36% female, 64% male; *T. brasiliensis*: 40% female, 60% male) and all reproductive groups (*M. velifer*: 76% non-reproductive, 24% reproductive; *T. brasiliensis*: 96% non-reproductive, 4% reproductive) were included in this study; all bats were adults. We determined our sample size (n=25 paired samples per species; n=50 paired samples total) using a two-sided, two-sample power analysis, assuming 92% power.

### Sample Collection

We collected paired blood from both the propatagial and intrafemoral veins per each individual bat (**Figure 1**). Before bleeding, veins were sterilized with 70% isopropyl and lanced with a sterile 26G needle, then blood was directly collected in a heparinized capillary tube. We used approximately 2 μL blood to make a thin blood smear on a sterile microscope slide, and the remaining blood was immediately transferred to a sterile 0.2 mL PCR tube to be used for total RBC, WBC, and RET counts. Because consecutive bleeds and relatively longer lags between animal capture and bleeding (e.g., bats bled later in the night) can induce physiological stress and affect cellular immunity (Davis, 2005; Davis et al., 2008), we recorded bleeding order from the propatagial and intrafemoral veins as well as the time between bat capture and blood collection. We aimed to collect an even ratio of bleeding order to reduce potential bias, and bats were on average held for 3.7 hours prior to blood collection.

Handling and care of bats were followed as described in Guidelines of the American Society of Mammalogists for the use of wild mammals in research (Sikes & Gannon 2011). All work was approved under University of Oklahoma Institutional Animal Care and Use Committee (IACUC) protocol 2022-0198 and scientific collection permit 10567389 from the Oklahoma Department of Wildlife Conservation. Proper personal protective equipment, KN95 masks, and leather and nitrile gloves were worn while handling and collecting samples to limit pathogen spread to or from bats (Olival et al., 2020).

### Hematology Measurements

To increase the accuracy of total RBC, WBC, and RET counts, we coupled blood smear analyses with Neubauer chamber counts (Fisher Scientific; depth 0.1 mm, area 0.0025 mm^2^). Blood was diluted 1:100 in a NaCl/EosinY solution for RBC counts and 1:1200 in a NaCl/Crystal Violet solution for WBC and RET counts (**Supplementary Table S1; Supplementary Table S2**). Sodium chloride and dye stock solutions were made no longer than one day before use, and dye stocks were stored in dark vessels. For consecutive nights of sampling in the field, stock solutions were stored in 4°C but not reused more than once. Within 1–2 hours of blood collection, each sample was stained and read for WBC, RBC, and RET total counts. WBC counts were calculated by averaging the number of cells observed in the four large 1 × 1-mm squares on either corner of the chamber; this equaled one read (Hansen, 2000). RBC and RET counts were calculated by averaging the number of cells observed in the five smaller 0.2 × 0.2-mm squares within the large middle square of the chamber; each five square average count equaled one read (Hansen, 2000). Each cell measure was read under 400X magnification with a binocular microscope (Euromex EBS1152EPLI). We used the following equations to calculate cells per microliter (Hansen, 2000):

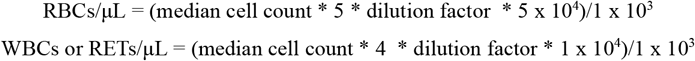

Blood smears were air dried and stored in sealed boxes with desiccant packs until they were fixed with methanol and stained with Wright–Giemsa (Quick III, Astral Diagnostics). To obtain WBC differentials, we identified up to 100 leukocytes in the monolayer of each blood smear under 1000X magnification (oil immersion) in a systematic grid pattern (Bain et al., 2006). Blood surface area of blood smears was not standardized, and some samples, despite having a large blood surface area, did not have 100 WBCs to identify. For downstream analyses, we categorized each blood smear by its surface area (small or large). All Neubauer chamber and blood smear measures were determined by one reader.

### Statistical Analysis

All analyses were performed in R software (R Core Team, 2023). To first assess consistency of each cell measure between venipuncture sites, we calculated intraclass correlation coefficients (ICCs) within each bat species using the *iccCounts* package (Carrasco, 2022). We classified levels of consistency as previously described: high (ICC ≥ 0.9), good (0.75 ≤ ICC ≤ 0.9), moderate (0.5 ≤ ICC ≤ 0.75) and poor (ICC < 0.5) (Haghayegh et al., 2020). We next used generalized linear mixed models (GLMMs) fit using the *lme4* package (Zuur, 2009) to test fixed effects of venipuncture site, bat species, and their interaction on all cell counts. All GLMMs included a random intercept for bat identification number and a random slope of venipuncture site. For total RBC, WBC, and RET counts, we used the median value across each read per sample; RBC and RET counts were modeled as negative binomial responses, while WBC counts were analyzed with a Poisson response, based on relationships between the cell count means and variance. WBC differential counts were modeled as binomial responses, considering the counts of each cell relative to that of all other cells identified; this approach allowed us to account for variation in total number of cells identified. We conducted post-hoc analyses and obtained marginal means from each GLMM using the *emmeans* package (Lenth et al., 2018). We then calculated the contrast percentage of the marginal mean values between veins within each species.

Though aiming to obtain an even ratio of vein bleeding order within our sample size goal (n=30 paired samples per species), our final ratio for both species was slightly uneven: 10 intrafemoral and 40 propatagial samples for *M. velifer*, 20 intrafemoral and 30 propatagial samples for *T. brasiliensis*. To assess the potential bias of bleeding order ratio, we excluded data from the first sampling trip (September 2023) to obtain a more even ratio: 10 intrafemoral and 30 propatagial samples for *M. velifer*, 20 intrafemoral and 20 propatagial samples for *T. brasiliensis*. Likewise, we ran additional sensitivity analyses that independently included blood smear size, vein bleeding order, and time between capture and blood collection as precision covariates; all covariates could not be included in the same GLMM due to sample size.

## Results

When assessing concordance in cell counts between propatagial and intrafemoral veins for *M. velifer*, consistency was high for RETs, basophils, eosinophils, segmented neutrophils, and total neutrophils; good for WBCs, banded neutrophils, and lymphocytes; moderate for monocytes; and poor for RBCs and NL ratios (**Supplementary Figure 1**). Similarly, for *T. brasiliensis*, RETs were highly consistent while eosinophils, segmented neutrophils, lymphocytes, and total neutrophils had only good consistency. Monocytes again were moderately consistent, and RBCs again had poor consistency. In contrast to *M. velifer*, WBCs, basophils, and banded neutrophils had poor consistency (**Supplementary Figure 1**).

Predicted means and confidence intervals of each cell measurement from our GLMMs are shown in **Figure 2** and **Supplementary Figure 2**. Across all our hematology values, our GLMMs found no significant effects of venipuncture site within or across our two bat species (**Supplementary Tables S3 & S4**). However, we did see species-level differences in WBC counts, RET counts, neutrophils, lymphocytes, NL ratios, segmented neutrophils, eosinophils, and basophils (**Supplementary Tables S3 & S4**). Results were consistent when analyzing only the subset of our data with a more even ratio of vein bleeding order (**Supplemental Table S5**). Similarly, results were unaffected when blood smear size, vein bleeding order, and time between capture and blood collection were independently accounted for in our GLMMs (**Supplemental Tables S6, S7, and S8**).

**Figure 2.**
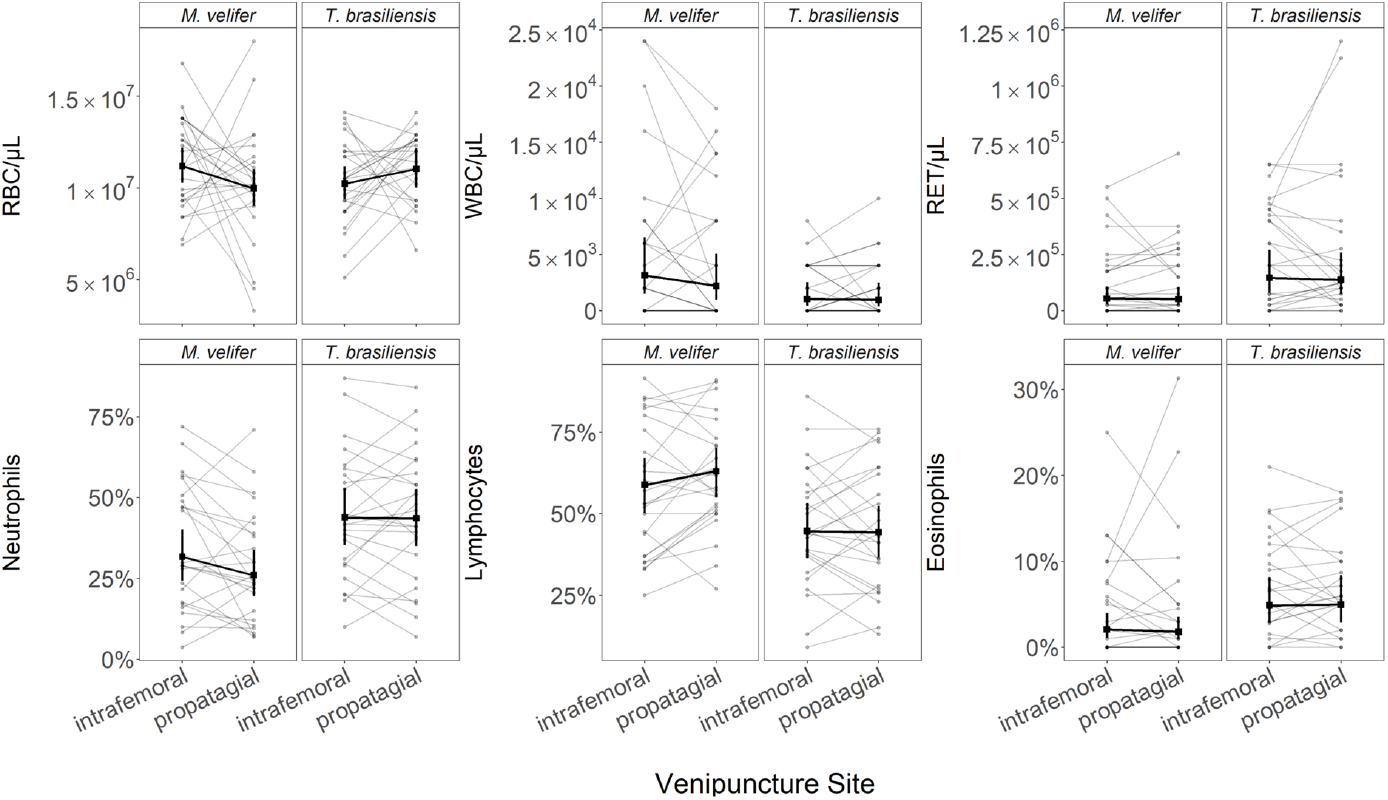
Hematology values as a function of venipuncture site, stratified by bat species. Paired vein data for an individual bat are shown through line segments. Bold coloring indicates the predicted means and 95% confidence intervals for each vein per species from our GLMMs. Model results are provided in Supplementary Table S3 for effects of venipuncture site, bat species, and their interaction.

Using marginal means from our GLMMs, we observed the largest inter-vein difference for the NL ratio in *T. brasiliensis*, where NL ratios from the propotagial vein were 67% higher than those from the intrafemoral vein (**Supplementary Figure 2**). For all other cell measurements, our GLMMs identified very small inter-vein differences: RBCs (2.7%); RETs, neutrophils, and lymphocytes (0.3%); monocytes and segmented neutrophils (0.2%); banded neutrophils (0.11%); eosinophils (0.08%); basophils (0.05%); andWBCs (0.02%). We observed similarly small inter-vein differences in *M. velifer*: RBCs (4.1%), monocytes (0.5%), banded neutrophils (0.42%), eosinophils (0.3%), WBCs (0.23%), basophils (0.2%), and RETs (0.1%). In comparison to *T. brasiliensis*, however, *M. velifer* had a higher percent contrast for mean cell counts between veins for neutrophils (5.7%), segmented neutrophils (4.7%), and lymphocytes (4%), but this species had a lower percent contrast for NL ratios (1.6%).

## Discussion

Standardized sampling protocols and proper methods reporting is important to facilitate robust comparisons in the fields of ecoimmunology and disease ecology. Our findings emphasize the relevance of these points for studies of wild bats, due to the high frequency of hematological techniques used to infer health status and infection prevalence in research on chiroptera and emerging infectious diseases (DeAnglis et al., 2024; Kovalchuk et al., 2017; Villalba-Alemán et al., 2020). Here, we tested the effect of two of the most common venipuncture sites in bats, the propatagial and intrafemoral veins, on multiple hematology measures of two tractable temperate bat systems in the Americas. We found that most cellular measures were highly repeatable, and all our GLMMs found no effect of venipuncture site on any of our hematology parameters. By contrast, we did find significant differences between bat species for many cell measures. We chose our two sample species (*T. brasiliensis* and *M. velifer*) not only based on economical and practical feasibility, as they are involved in long-term projects on bat immunity and infection (Becker et al. 2025), but also to capture variability within two of the largest and globally distributed chiropteran families, the Molossidae (*T. brasiliensis*) and Vespertilionidae (*M. velifer*). As aforementioned, *T. brasiliensis* and *M. velifer* vary in multiple ecological traits, such as their life-history strategy (i.e., long-distance migration and lack of hibernation versus resident or short-distance migratory status and hibernation). The significant differences detected among multiple cell measures between these two species could thus be attributed to their unique wintering strategies (Auteri, 2022). Furthermore, we are uncertain what could explain the noteworthy difference in the estimated means of the neutrophil and lymphocyte counts by venipuncture site between *T. brasiliensis* and *M. velifer*. These results could be another cause of taxonomic family or life-history strategy differences. Future comparisons of cellular immunity on more species representatives within Molossidae and Vespertilionidae, as well as targeted seasonal sampling, could address these observed differences.

Based on the ubiquitous lack of venipuncture effects across hematology measurements, our findings suggest little to no differences in cellular pathways or composition throughout the body of *T. brasiliensis* or *M. velifer*. Previous studies in species other than bats suggest that lymph hemodilution, lymph contamination (Crooks et al., 2023; Neiffer et al., 2021; Rodríguez-Almonacid et al., 2022), hemolysis, and sera dilution (Comazzi et al., 2023) could explain different hematology values at each vein site due to their proximity to lymphatics. Other work has suggested that microanatomical obstacles in the circulatory system could explain observed hematological differences by venipuncture site (Mylniczenko et al., 2006), and our null findings imply that such effects may not be present among these sympatric but phylogenetically distinct species

When conducting our comparisons across venipuncture site within and across bat species, we aimed to eliminate as many biases as possible, including statistically powerful and equal sample-size groups for each species, ensuring identical methods were carried out for each blood draw, and having only one reader for all hematology metrics. We also considered bleeding order of the veins, time between capture and bleeding, and differences in blood smear size, with results consistent after statistically adjusting for these variables. However, we acknowledge some limitations still remain in these data, such as not having an equal bleeding order ratio in the final dataset, sampling occurring during different months of the year, variation in handler and bleeder, and variability in the time between blood collection and Neubauer chamber readings.

Given our focus on two bat species within two families, for two commonly used veins in blood collection, we encourage testing of venipuncture sites on other veins used in bat surveys, testing between age, reproductive status, and sex groups, and expanding these tests to other bat species (and families) to further support the non-significant effects observed here. We also encourage testing effects of venipuncture site on health metrics other than hematology that are increasingly used in bat immunology, such as gene expression and protein abundance data afforded through transcriptomics and proteomics (Arnold et al., 2018; Becker & Banerjee, 2023). Future studies testing venipuncture sites in bats should also consider the limitations we note above. Any significant findings of future studies could be used to determine a correction factor between vein types, which could then be applied more generally to ensure comparisons of health metrics in bats are robust and appropriately assessed.

Lastly, our results should provide some reassurance for the fields of ecoimmunology and disease ecology, as our findings suggest negligible effects of blood collection from either the propatagial or intrafemoral vein on typical hematology measurements. Because of this, we believe there should be negligible miscalculation or interpretation of results from previous hematology studies with molossid or vesper bat species that collected peripheral blood from a combination of the propatagial and intrafemoral veins or for comparative analyses where different studies use one or the other vein for blood collection. These findings are also valuable for future similar intra- and inter-species comparative analyses, as vein type would not be a necessary precision covariate (Becker et al. 2025). However, we suggest that vein type should still be consistently reported in publications, and it could still be valuable to establish baseline parameter ranges by each vein site (Neiffer et al., 2021). For species of conservation concern, researchers could use our findings to motivate switching vein types for blood collection during sample collection to obtain the minimum blood requirements for assays to inform health status (Ohmer et al., 2021).

## Acknowledgements

This project was funded by the National Science Foundation (DBI 2515340), Edward Mallinckrodt, Jr. Foundation, and the American Society of Mammalogists (Grants-in-Aid of Research Award). Fieldwork was made possible through collaboration with Oklahoma State Parks, the Oklahoma Department of Wildlife Conservation, the University of Central Oklahoma, and the Selman Living Laboratory. We acknowledge Jadeyn Lindsey and Marguerite Hall for significant contributions to field sampling. We thank Mika O’Shea for illustrating the bat shown in Figure 1 and Jon Alonzo and Ralph Simon for providing the venipuncture site photographs for Figure 1A-B. We thank Meagan Allira, Kristin Dyer, Caroline Cummings, Lauren Lock, and Mackenzie Hightower for providing additional fieldwork support. Lastly, we thank Emily Gallichotte for contributing to manuscript revision.

## References

Alharbi, A. H. V A. C.,, Lin, M., Ashwini, B., Jabarulla, M. Y., & Shah, M. A. (2022). Detection of Peripheral Malarial Parasites in Blood Smears Using Deep Learning Models. Computational Intelligence and Neuroscience, 2022, 3922763.

Ammerman, L. K., Hice, C. L., & Schmidly, D. J. (2012). Bats of Texas. Texas A & M University Press.

Arnold, C. E., Guito, J. C., Altamura, L. A., Lovett, S. P., Nagle, E. R., Palacios, G. F., Sanchez-Lockhart, M., & Towner, J. S. (2018). Transcriptomics reveal antiviral gene induction in the Egyptian rousette bat is antagonized in vitro by Marburg virus infection. Viruses, 10(11), 607.

Auteri, G. G. (2022). A conceptual framework to integrate cold-survival strategies: torpor, resistance and seasonal migration. Biology Letters, 18(5), 20220050.

Ayala-Berdon, J., & Solis-Cardenas, V. (2017). New record and site characterization of a hibernating colony of Myotis velifer in a mountain ecosystem of central Mexico. Therya, 8(2), 175.

Bain, B. J., Lewis, S. M., & Bates, I. (2006). Basic haematological techniques. Dacie and Lewis Practical Haematology, 4, 19–46.

Becker, D. J., & Banerjee, A. (2023). Coupling field and laboratory studies of immunity and infection in zoonotic hosts. The Lancet. Microbe, 4(5), e285–e287.

Becker, D. J., Dyer, K. E., Lock, L. R., Olbrys, B. L., Pladas, S. A., Sukhadia, A. A., Demory, B., Nunes Batista, J. M., Pineda, M., Simmons, N. B., Adams, A. M., Frick, W. F., O’Mara, M. T., & Volokhov, D. V. (2025). Geographically widespread and novel hemotropic mycoplasmas and bartonellae in Mexican free-tailed bats and sympatric North American bat species. mSphere, 10(1), e0011624.

Becker, D. J., Vicente-Santos, A., Reers, A., Ansil, B. R., O’Shea, M., Cummings, C., Roistacher, A., Quintela-Tizon, R., Pereira, M., Rosen, J., Banerjee, A., & Frank, H. (2025). Diverse hosts, diverse immune systems: evolutionary variation in bat immunology. EcoEcoRxiv. 10.32942/X2HD0D

Betke, B. A., Gottdenker, N. L., Meyers, L. A., & Becker, D. J. (2024). Ecological and evolutionary characteristics of anthropogenic roosting ability in bats of the world. iScience, 27(7), 110369.

Betke, M., Hirsh, D. E., Makris, N. C., McCracken, G. F., & Cleveland, C. J. (2008). Thermal Imaging Reveals Significantly Smaller Brazilian Free-Tailed Bat Colonies Than Previously Estimated. Journal of Mammalogy, 89(1), 18–24.

Bryant, A. (2013). From “Bat Caves” to “Alabaster Caverns” a History of the Use and Conservation of Alabaster Caverns State Park. ProQuest.

Burrell, G. E., & Bergeson, S. (2022). Roosting behavior of northern long-eared bats (Myotis septentrionalis) in an urban-adjacent forest fragment. Forests. 10.3390/f13121972

Caire, W., & Loucks, L. S. (2010). Loss in Mass by Hibernating Cave Myotis, Myotis velifer (Chiroptera: Vespertilionidae) in Western Oklahoma. The Southwestern Naturalist, 55(3), 323–330.

Caire, W., Smith, J. F., McGuire, S., & Royce, M. A. (1984). Early foraging behavior of insectivorous bats in western Oklahoma. Journal of Mammalogy, 65, 319–324.

Carrasco, J. L. (2022). IccCounts: An R package to estimate the intraclass correlation coefficient for assessing agreement with count data. The R Journal. 10.32614/rj-2022-034

Cheng, T. L., Reichard, J. D., Coleman, J. T. H., Weller, T. J., Thogmartin, W. E., Reichert, B. E., Bennett, A. B., Broders, H. G., Campbell, J., Etchison, K., Feller, D. J., Geboy, R., Hemberger, T., Herzog, C., Hicks, A. C., Houghton, S., Humber, J., Kath, J. A., King, R. A., … Frick, W. F. (2021). The scope and severity of white-nose syndrome on hibernating bats in North America. Conservation Biology: The Journal of the Society for Conservation Biology, 35(5), 1586–1597.

Comazzi, S., Guanziroli, S., Giordano, A., Formenti, N., Trogu, T., Corlatti, L., Luzzago, C., & Ferrari, N. (2023). Comparison between cavernous sinus and jugular vein as post mortem sampling sites for blood metabolic profiles in wild red deer (Cervus elaphus). Zeitschrift Für Jagdwissenschaft, 69. 10.1007/s10344-023-01715-w

Cornelius Ruhs, E., Becker, D. J., Oakey, S. J., Ogunsina, O., Fenton, M. B., Simmons, N. B., Martin, L. B., & Downs, C. J. (2021). Body size affects immune cell proportions in birds and non-volant mammals, but not bats. The Journal of Experimental Biology, 224(13). 10.1242/jeb.241109

Crooks, G. C., Calle, P. P., Moore, R. P., McClave, C., Toledo, P., Gomez, N. A., Perez, V. B., Tewfik, A., Rao, S., & Sadar, M. J. (2023). HEMATOLOGIC AND BIOCHEMICAL VALUES OF FREE-RANGING HAWKSBILL SEA TURTLES (ERETMOCHELYS IMBRICATA) IN GLOVER’S REEF, BELIZE. Journal of Zoo and Wildlife Medicine: Official Publication of the American Association of Zoo Veterinarians, 54(1), 49–55.

Danielson, J. R., Williams, J. A., Sherwin, R. E., Ekholm, K. L., & Hamilton, B. T. (2022). Seasonal variation in age, sex, and reproductive status of Mexican free-tailed bats. Population Ecology, 64(3), 254–266.

Davis, A. K. (2005). Effect of handling time and repeated sampling on avian white blood cell counts. Journal of Field Ornithology, 76(4), 334–338.

Davis, A. K., Maney, D. L., & Maerz, J. C. (2008). The use of leukocyte profiles to measure stress in vertebrates: a review for ecologists. Functional Ecology, 22(5), 760–772.

DeAnglis, I. K., Andrews, B. R., Lock, L. R., Dyer, K. E., Yang, A., Volokhov, D. V., Fenton, M. B., Simmons, N. B., Downs, C. J., & Becker, D. J. (2024). Bat cellular immunity varies by year and dietary habit amidst land conversion. Conservation Physiology, 12(1), coad102.

Donald W. Tinkle, I.G.P. (1965). A study of hibernating populations of Myotis velifer in northwestern Texas. Journal of Mammalogy, 46(4), 612–633.

Dziki-Michalska, K., Tajchman, K., Kowalik, S., & Wójcik, M. (2024). The levels of cortisol and selected biochemical parameters in Red Deer harvested during stalking hunts. Animals: An Open Access Journal from MDPI, 14. 10.3390/ani14071108

Eshar, D., Gancz, A. Y., Avni-Magen, N., Wagshal, E., Pohlman, L. M., & Mitchell, M. A. (2016). SELECTED PLASMA BIOCHEMISTRY ANALYTES OF HEALTHY CAPTIVE SULCATA (AFRICAN SPURRED) TORTOISES (CENTROCHELYS SULCATA). Journal of Zoo and Wildlife Medicine: Official Publication of the American Association of Zoo Veterinarians, 47(4), 993–999.

Eshar, D., & Weinberg, M. (2010). Venipuncture in bats. Lab Animal, 39(6), 175–176.

Fitch, J. H., Shump, K. A., join(‘‘., & Shump, A. U. (1981). Myotis velifer. Mammalian Species, 149, 1.

Gottdenker, N. L., & Jacobson, E. R. (1995). Effect of venipuncture sites on hematologic and clinical biochemical values in desert tortoises (Gopherus agassizii). American Journal of Veterinary Research, 56(1), 19–21.

Gregory, T. M., Hubbard, C., Schlake, E., Mejia, D., Passingham, K. R., Lewbart, G. A., & Harrison, T. M. (2022). EVALUATION OF PROGNOSTIC INDICATORS FOR INJURED TURTLES PRESENTING TO A WILDLIFE CLINIC. Journal of Zoo and Wildlife Medicine: Official Publication of the American Association of Zoo Veterinarians, 53(1), 209–213.

Gupta, N., & Nigar, S. (2020). Detection of blood parasites and estimation of hematological indices in fish. In Springer Protocols Handbooks (pp. 43–73). Springer US.

Haghayegh, S., Kang, H.-A., Khoshnevis, S., Smolensky, M. H., & Diller, K. R. (2020). A comprehensive guideline for Bland-Altman and intra class correlation calculations to properly compare two methods of measurement and interpret findings. Physiological Measurement, 41(5), 055012.

Hansen, P. J. (2000). Use of a Hemacytometer. Department of Animal Science, University of Florida, 10–15.

Humphrey, S. R., & Oli, M. K. (2015). Population dynamics and site fidelity of the cave bat,Myotis velifer, in Oklahoma. Journal of Mammalogy, 96(5), 946–956.

Irving, A. T., Ahn, M., Goh, G., Anderson, D. E., & Wang, L.-F. (2021). Lessons from the host defences of bats, a unique viral reservoir. Nature, 589(7842), 363–370.

Iskali, G., & Zhang, Y. (2015). Guano subsidy and the invertebrate community in bracken cave: The world’s larges colony of bats. Journal of Cave and Karst Studies: The National Speleological Society Bulletin, 77(1), 28–36.

Kern, M. D., & De Graw, W. A. (1978). Differences in the Composition of Venous and Cardiac Blood from White-Crowned Sparrows. The Condor, 80(2), 230–234.

Kloskowski, J., Kaczanowska, E., Krogulec, J., & Grela, P. (2017). Hematological indicators of habitat quality: Erythrocyte parameters reflect greater parental effort of Red-necked Grebes under ecological trap conditions. The Condor, 119(2), 239–250.

Kovalchuk, L., Mishchenko, V., Chernaya, L., Snitko, V., & Mikshevich, N. (2017). Haematological parameters of pond bats (Myotis dasycneme Boie, 1825 Chiroptera: Vespertilionidae) in the Ural Mountains. Zoology and Ecology, 27(2), 168–175.

Krutzsch, P. H. (2009). The Reproductive Biology of the Cave Myotis (Myotis velifer). Acta Chiropterologica, 11(1), 89–104.

Lenth, R., Singmann, H., Love, J., Buerkner, P., & Herve, M. (2018). Emmeans: Estimated marginal means. AKA Least-Squares Means, 1(7). https://scholar.google.com/citations?user=howgU0gAAAAJ&hl=en&oi=sra

López-Olvera, J. R., Montané, J., Marco, I., Martínez-Silvestre, A., Soler, J., & Lavín, S. (2003). EFFECT OF VENIPUNCTURE SITE ON HEMATOLOGIC AND SERUM BIOCHEMICAL PARAMETERS IN MARGINATED TORTOISE. Journal of Wildlife Diseases, 39(4), 830–836.

Maceda-Veiga, A., Figuerola, J., Martínez-Silvestre, A., Viscor, G., Ferrari, N., & Pacheco, M. (2015). Inside the Redbox: applications of haematology in wildlife monitoring and ecosystem health assessment. The Science of the Total Environment, 514, 322–332.

McCracken, G. F., Bernard, R. F., Gamba-Rios, M., Wolfe, R., Krauel, J. J., Jones, D. N., Russell, A. L., & Brown, V. A. (2018). Rapid range expansion of the Brazilian free-tailed bat in the southeastern United States, 2008–2016. Journal of Mammalogy, 99(2), 312–320.

Mylniczenko, N. D., Curtis, E. W., Wilborn, R. E., & Young, F. A. (2006). Differences in hematocrit of blood samples obtained from two venipuncture sites in sharks. American Journal of Veterinary Research, 67(11), 1861–1864.

Neiffer, D. L., Hayek, L.-A. C., Conyers, D., Daneault, A., Sincage, J., & Yawn, J. (2021). COMPARISON OF SUBCARAPACIAL SINUS AND BRACHIAL VEIN PHLEBOTOMY SITES FOR BLOOD COLLECTION IN FREE-RANGING GOPHER TORTOISES (GOPHERUS POLYPHEMUS). Journal of Zoo and Wildlife Medicine: Official Publication of the American Association of Zoo Veterinarians, 52(3), 966–974.

Ohmer, M. E. B., Costantini, D., Czirják, G.Á., Downs, C. J., Ferguson, L. V., Flies, A., Franklin, C. E., Kayigwe, A. N., Knutie, S., Richards-Zawacki, C. L., & Cramp, R. L. (2021). Applied ecoimmunology: using immunological tools to improve conservation efforts in a changing world. Conservation Physiology, 9(1), coab074.

Olival, K. J., Cryan, P. M., Amman, B. R., Baric, R. S., Blehert, D. S., Brook, C. E., Calisher, C. H., Castle, K. T., Coleman, J. T. H., Daszak, P., Epstein, J. H., Field, H., Frick, W. F., Gilbert, A. T., Hayman, D. T. S., Ip, H. S., Karesh, W. B., Johnson, C. K., Kading, R. C., … Wang, L.-F. (2020). Possibility for reverse zoonotic transmission of SARS-CoV-2 to free-ranging wildlife: A case study of bats. PLoS Pathogens, 16(9), e1008758.

Pouletty, N. (2010). Blood smears: evaluation of thrombocytes and leukocytes. Point Vétérinaire. https://www.cabdirect.org/cabdirect/abstract/20103142455

R Core Team. (2023). R: A language and environment for statistical computing. https://www.R-project.org/

Rodríguez-Almonacid, C. C., Vargas-León, C. M., Moreno-Torres, C. A., & Matta Camacho, N. E. (2022). Consideraciones para la obtención de sangre en tortugas: sitios de venopunción y anticoagulantes. Revista MVZ Cordoba, 27(2), e2256.

Ruoss, S., Becker, N. I., Otto, M. S., Czirják, G.Á., & Encarnação, J. A. (2019). Effect of sex and reproductive status on the immunity of the temperate bat Myotis daubentonii. Zeitschrift Für Saugetierkunde [Mammalian Biology], 94(1), 120–126.

Santana, S. E., Dial, T. O., Eiting, T. P., & Alfaro, M. E. (2011). Roosting ecology and the evolution of pelage markings in bats. PloS One, 6(10), e25845.

Schinnerl, M., Aydinonat, D., Schwarzenberger, F., & Voigt, C. C. (2011). Hematological survey of common neotropical bat species from Costa Rica. Journal of Zoo and Wildlife Medicine: Official Publication of the American Association of Zoo Veterinarians, 42(3), 382–391.

Schneeberger, K., Czirják, G.Á., & Voigt, C. C. (2013). Measures of the constitutive immune system are linked to diet and roosting habits of neotropical bats. PloS One, 8(1), e54023.

Sheldon, L. D., Chin, E. H., Gill, S. A., Schmaltz, G., Newman, A. E. M., & Soma, K. K. (2008). Effects of blood collection on wild birds: an update. Journal of Avian Biology, 39(4), 369–378.

Sikes, R. S. (2016). 2016 Guidelines of the American Society of Mammalogists for the use of wild mammals in research and education. Journal of Mammalogy, 97(3), 663–688.

Tovstukha, I., Fritze, M., Kravchenko, K., & Kovalov, V. (2024). https://www.researchgate.net/profile/Marcus-Fritze-2/publication/377402470_Pilot_study_suggests_cellular_immunity_changes_in_bats_from_urban_landscapes/links/65a51d8a8ee032139ae7bce0/Pilot-study-suggests-cellular-immunity-changes-in-bats-from-urban-landscapes.pdf

Villa, B. R., & Lendell Cockrum, E. (1962). Migration in the Guano Bat Tadarida Brasiliensis Mexicana (Saussure). Journal of Mammalogy, 43(1), 43–64.

Villalba-Alemán, E., Bustos, X., Crisante, G., de Jesús, R., Mata, J., Pereira, F., & Muñoz-Romo, M. (2020). Hematological characterization of common bats in urban areas from Mérida (Venezuela), and observations on possible hemopathogens. Acta Chiropterologica, 22(2). 10.3161/15081109acc2020.22.2.017

Wiederholt, R., López-Hoffman, L., Svancara, C., McCracken, G., Thogmartin, W., Diffendorfer, J. E., Mattson, B., Bagstad, K., Cryan, P., Russell, A., Semmens, D., & Medellín, R. A. (2015). Optimizing conservation strategies for Mexican free-tailed bats: a population viability and ecosystem services approach. Biodiversity and Conservation, 24(1), 63–82.

Wilkins, K. T. (1989). Tadarida brasiliensis. Mammalian Species, 331, 1.

William, C., & Samanie, L. L. (2010). Loss in Mass by Hibernating Cave Myotis, Myotis velifer (Chiroptera: Vespertilionidae) in Western Oklahoma. The Southwestern Naturalist, 55(3), 323–330.

Wolfensohn, S., & Lloyd, M. (2013). Handbook of laboratory animal management and welfare (4th ed.) [PDF]. Wiley-Blackwell.

Zuur, A. F. (2009). Mixed effects models and extensions in ecology with R [PDF]. Springer. 10.1007/978-0-387-87458-6

