## Supplemental Materials for "The effect of venipuncture site on hematology of bats: implications for comparative analyses"

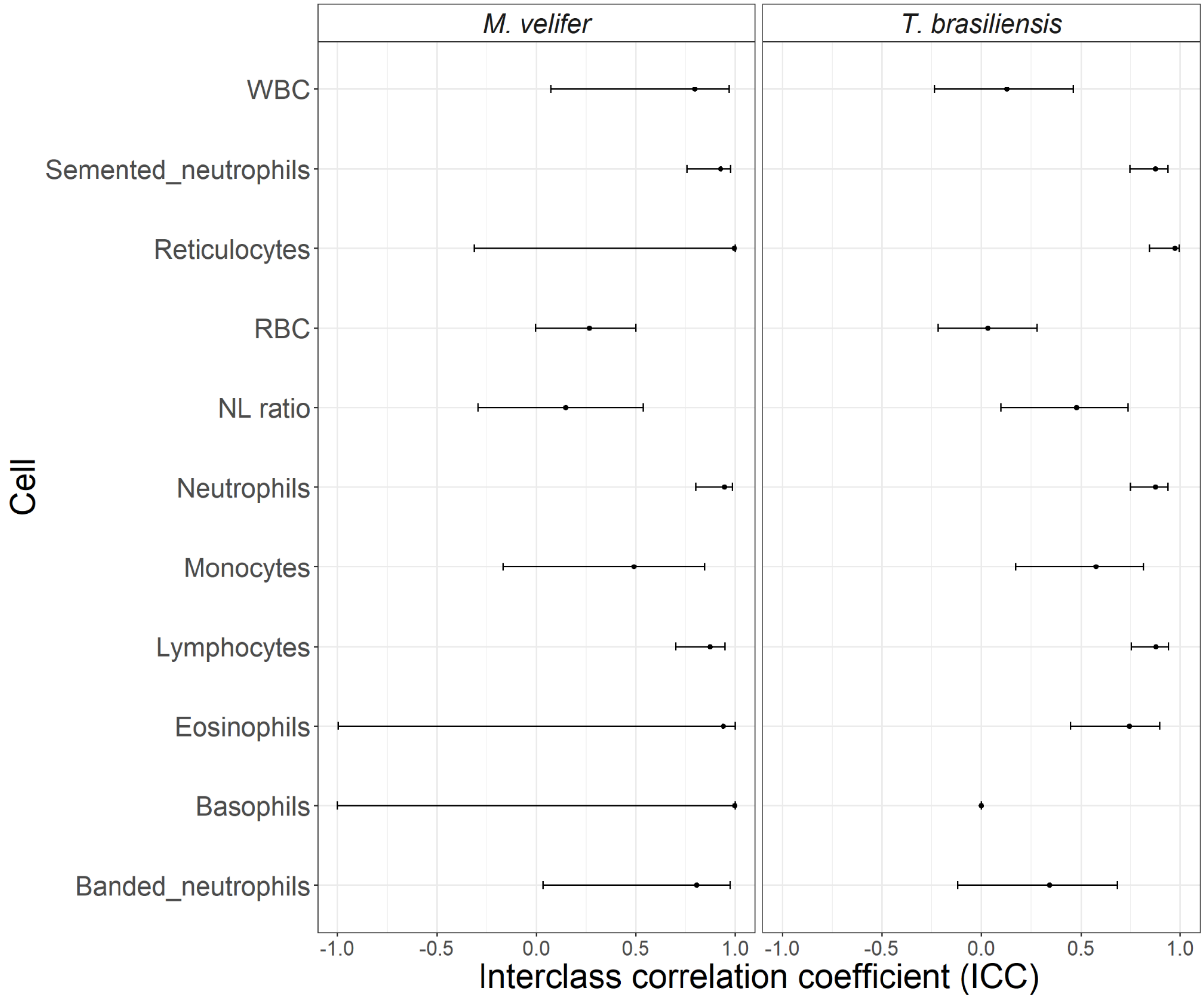


**Figure S1:** Repeatability estimates and 95% confidence intervals for each cell measure. Where confidence intervals are not present, values were exponentially negative.

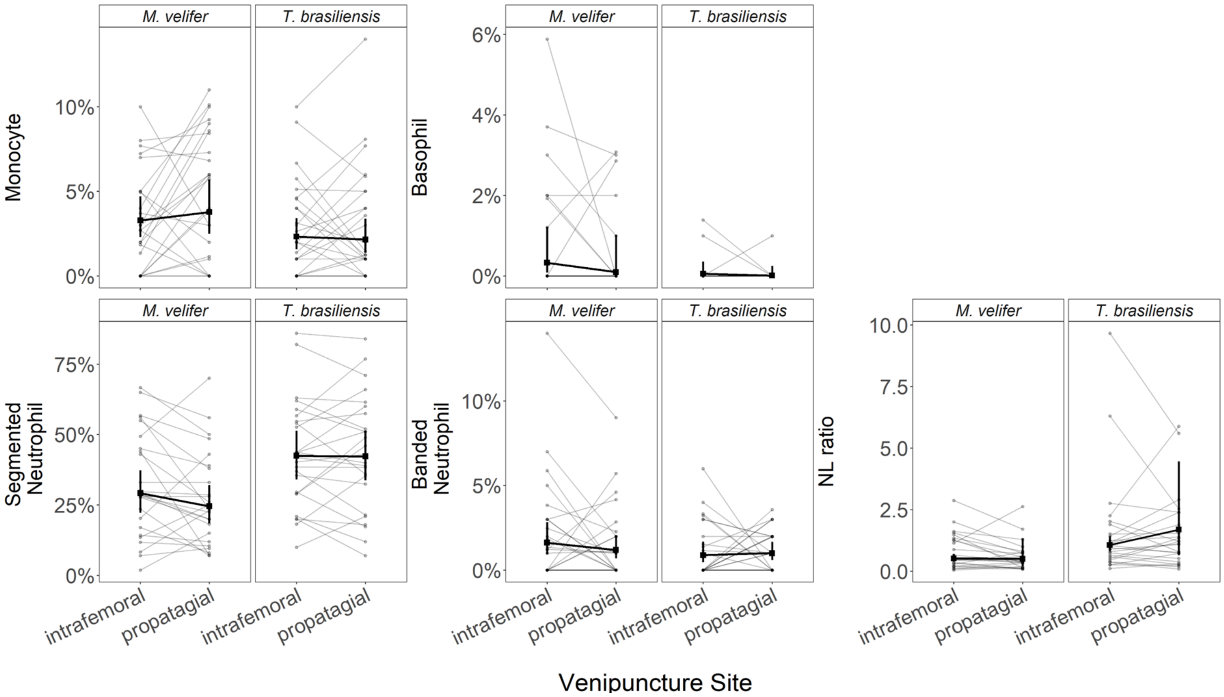


**Figure S2:** Hematology values as a function of venipuncture site, stratified by bat species. Paired vein data for an individual bat are shown through line segments. Bold coloring indicates the predicted means and 95% confidence intervals for each vein per species from our GLMMs. Model results Summary statistical values are provided in Supplementary Table S4 for effects of venipuncture site, bat species, and their interaction.

| **Table S1**: Formulas for Neuber chamber dye solutions. All volumes provided in µL units. Blood/dye solutions were mixed in 2mL vessels. Blood was added within 5 minutes of counting, the vessel is gently inverted to sufficiently mix, and then allowed to sit for ~2 minutes to allow cells to absorb the stain. Blood/dye solutions are loaded into the Nauber chamber and let it sit for another ~1-2 minutes to allow the cells to evenly disperse in the chamber. | | | | |
|---|---|---|---|---|
| Cell Type | Blood | 0.75% sodium chloride in mH20 | 0.5% Eosin Y | 0.5% Crystal Violet |
| RBC | 1 | 1,175 | 24 | — |
| WBC | 6 | 582 | — | 12 |
| RET | 6 | 582 | — | 12 |

| **Table S2:** Formulas for 10mL stock solutions of Neuber chamber dyes. Units per reagent is provided within each cell. | | | | | | |
|---|---|---|---|---|---|---|
| Solution | 95% EtOH | Glacial acetic acid | MeOH | mH20 | Eosin Y | Crystal Violet |
| 0.5% EosinY, 0.1% glacial acetic acid, 20% v/v EtOH | 2 mL | 10 µL | — | 8 mL | 0.05 g | — |
| 0.5% Crystal violet, 20% v/v MeOH | — | — | 2 mL | 8 mL | — | 0.05 g |

**Table S3**: Summary of GLMM results for cell measures included in Figure 2, using type II ANOVA tests .

| Cell Measure | Variable | $\chi^2$ | df | *p* |
|---|---|---|---|---|
| RBC | Vein | 0.1402 | 1 | 0.70811 |
| | Species | 0.0112 | 1 | 0.91569 |
| | Vein:Species | 3.3425 | 1 | 0.06751 |
| WBC | Vein | 0.4268 | 1 | 0.51355 |
| | Species | **4.8307** | **1** | **0.02796*** |
| | Vein:Species | 0.3482 | 1 | 0.55514 |
| RET | Vein | 0.1196 | 1 | 0.72944 |
| | Species | **5.0185** | **1** | **0.02508*** |
| | Vein:Species | 0.0006 | 1 | 0.98037 |
| Neutrophils | Vein | 1.8590 | 1 | 0.172739 |
| | Species | **7.0288** | **1** | **0.008021**** |
| | Vein:Species | 1.8184 | 1 | 0.177506 |
| Lymphocytes | Vein | 0.8175 | 1 | 0.36591 |
| | Species | **9.4342** | **1** | **0.00213**** |
| | Vein:Species | 1.2047 | 1 | 0.27238 |
| Eosinophils | Vein | 0.0106 | 1 | 0.91813 |
| | Species | **5.6145** | **1** | **0.01781*** |
| | Vein:Species | 0.2450 | 1 | 0.62059 |

**Table S4**: Summary of GLMM results for cell measures included in Supplementary Figure 2, using type II ANOVA tests.

| Cell Type | Variable | $\chi^2$ | df | *p* |
|---|---|---|---|---|
| Monocyte | Vein | 0.0687 | 1 | 0.79324 |
| | Species | 3.4638 | 1 | 0.06272 |
| | Vein:Species | 0.5042 | 1 | 0.47766 |
| Basophil | Vein | 1.0455 | 1 | 0.30654 |
| | **Species** | **5.1199** | **1** | **0.02365*** |
| | Vein:Species | 0.1186 | 1 | 0.73055 |
| Segmented neutrophil | Vein | 1.3869 | 1 | 0.238937 |
| | **Species** | **8.0534** | **1** | **0.004542**** |
| | Vein:Species | 1.3914 | 1 | 0.238164 |
| Banded neutrophil | Vein | 0.1448 | 1 | 0.7035 |
| | Species | 1.5605 | 1 | 0.2116 |
| | Vein:Species | 0.8798 | 1 | 0.3483 |
| NL ratio | Vein | 0.4100 | 1 | 0.5219486 |
| | **Species** | **12.0194** | **1** | **0.0005265***** |
| | Vein:Species | 0.5411 | 1 | 0.4619589 |

**Table S5:** Summary of GLMM results for cell measures fit to the subset of data with a more even ratio of vein bleeding order, using type II ANOVA tests.

| Cell Type | Variable | $\chi^2$ | df | *p* |
|---|---|---|---|---|
| RBC | Vein | 0.8953 | 1 | 0.3440 |
| | Species | 1.1201 | 1 | 0.2899 |
| | Vein:Species | 2.5823 | 1 | 0.1081 |
| WBC | Vein | 0.9244 | 1 | 0.3363 |
| | **Species** | **4.6806** | **1** | **0.0305*** |
| | Vein:Species | 0.3688 | 1 | 0.5437 |
| RET | Vein | 1.6569 | 1 | 0.198028 |
| | **Species** | **7.6330** | **1** | **0.005731**** |
| | Vein:Species | 0.0008 | 1 | 0.976954 |
| Neutrophils | Vein | 1.2152 | 1 | 0.2703025 |
| | **Species** | **14.4852** | **1** | **0.0001413***** |
| | Vein:Species | 2.3206 | 1 | 0.1276740 |
| Lymphocytes | Vein | 0.1556 | 1 | 0.6932 |
| | **Species** | **17.0460** | **1** | **$3.649 \times 10^{-5}$***** |
| | Vein:Species | 101438 | 1 | 0.2849 |
| Monocyte | Vein | 0.4396 | 1 | 0.50731 |
| | Species | 2.7862 | 1 | 0.09508 |
| | Vein:Species | 0.5640 | 1 | 0.45266 |
| Basophil | Vein | 0.7313 | 1 | 0.3925 |
| | Species | 2.1559 | 1 | 0.1420 |
| | Vein:Species | 0.0272 | 1 | 0.8690 |
| Eosinophils | Vein | 0.0538 | 1 | 0.81665 |
| | Species | 3.1564 | 1 | 0.07563 |
| | Vein:Species | 0.0258 | 1 | 0.87236 |

| | | | | |
|---|---|---|---|---|
| Banded neutrophil | Vein | 0.1796 | 1 | 0.6717 |
| | Species | 1.7425 | 1 | 0.1868 |
| | Vein:Species | 0.6061 | 1 | 0.4362 |
| Segmented neutrophil | Vein | 0.8802 | 1 | 0.3482 |
| | **Species** | **16.0933** | **1** | **$6.03 \times 10^{-5}$***** |
| | Vein:Species | 1.9286 | 1 | 0.1649 |
| NL ratio | Vein | 0.0191 | 1 | 0.890030 |
| | **Species** | **10.8103** | **1** | **0.001009**** |
| | Vein:Species | 0.4407 | 1 | 0.506797 |

| **Table S6:** Summary of GLMMs that include blood smear size, using type II ANOVA tests. | | | | |
|---|---|---|---|---|
| **Cell Type** | **Variable** | $\chi^2$ | **df** | ***p*** |
| Neutrophils | Vein | 0.8724 | 1 | 0.350284 |
| | **Species** | **6.8137** | **1** | **0.009046**** |
| | Slide Size | 1.0382 | 1 | 0.308244 |
| | Vein:Species | 1.7300 | 1 | 0.188417 |
| Lymphocytes | Vein | 0.6493 | 1 | 0.420370 |
| | **Species** | **9.4046** | **1** | **0.002164**** |
| | Slide Size | 0.0066 | 1 | 0.935230 |
| | Vein:Species | 1.1965 | 1 | 0.274014 |
| Monocyte | Vein | 0.0555 | 1 | 0.81376 |
| | Species | 3.4692 | 1 | 0.06252 |
| | Slide Size | 0.0005 | 1 | 0.98218 |
| | Vein:Species | 0.4974 | 1 | 0.48065 |
| Basophil | Vein | 1.2892 | 1 | 0.25619 |
| | **Species** | **5.1413** | **1** | **0.02336*** |
| | Slide Size | 0.6048 | 1 | 0.43675 |
| | Vein:Species | 0.1047 | 1 | 0.74628 |
| Eosinophils | Vein | 0.1686 | 1 | 0.68134 |
| | **Species** | **5.3966** | **1** | **0.02018*** |
| | Slide Size | 3.0050 | 1 | 0.08301 |
| | Vein:Species | 0.4068 | 1 | 0.52359 |
| Banded neutrophil | Vein | 0.1563 | 1 | 0.6926 |
| | Species | 1.5623 | 1 | 0.2113 |
| | Slide Size | 0.0360 | 1 | 0.8496 |
| | Vein:Species | 0.8860 | 1 | 0.3466 |

| | | | | |
|---|---|---|---|---|
| Segmented neutrophil | Vein | 0.5676 | 1 | 0.45123 |
| | **Species** | **7.8542** | **1** | **0.00507**** |
| | Slide Size | 1.0487 | 1 | 0.30580 |
| | Vein:Species | 1.3075 | 1 | 0.25285 |
| NL ratio | Vein | 0.0791 | 1 | 0.778515 |
| | **Species** | **7.8646** | **1** | **0.005041**** |
| | Slide Size | 0.0322 | 1 | 0.857580 |
| | Vein:Species | 0.7785 | 1 | 0.377608 |

| **Table S7:** Summary of GLMMs that include vein bleeding order, using type II ANOVA tests. | | | | |
|---|---|---|---|---|
| **Cell Type** | **Variable** | $\chi^2$ | **df** | ***p*** |
| Neutrophils | Vein | 1.8670 | 1 | 0.17182 |
| | **Species** | **5.8541** | **1** | **0.01554*** |
| | Bleeding order | 0.7882 | 1 | 0.37464 |
| | Vein:Species | 1.8013 | 1 | 0.17956 |
| Lymphocytes | Vein | 0.8387 | 1 | 0.35977 |
| | **Species** | **7.3040** | **1** | **0.00688**** |
| | **Bleeding order** | **4.1024** | **1** | **0.04282*** |
| | Vein:Species | 1.2146 | 1 | 0.27043 |
| Monocyte | Vein | 0.8908 | 1 | 0.345252 |
| | **Species** | **7.3487** | **1** | **0.006711**** |
| | **Bleeding order** | **9.8903** | **1** | **0.001662**** |
| | Vein:Species | 0.5120 | 1 | 0.474293 |
| Basophil | Vein | 1.7149 | 1 | 0.19035 |
| | **Species** | **6.3871** | **1** | **0.01150*** |
| | Bleeding order | 3.2471 | 1 | 0.07155 |
| | Vein:Species | 0.1864 | 1 | 0.66596 |
| Eosinophils | Vein | 0.0987 | 1 | 0.753370 |
| | **Species** | **3.8549** | **1** | **0.049601*** |
| | **Bleeding order** | **7.7626** | **1** | **0.005334**** |
| | Vein:Species | 0.2912 | 1 | 0.589456 |
| Banded neutrophil | Vein | 0.1367 | 1 | 0.7116 |
| | Species | 1.6916 | 1 | 0.1934 |
| | Bleeding order | 0.1730 | 1 | 0.6775 |
| | Vein:Species | 0.8792 | 1 | 0.3484 |

| | | | | |
|---|---|---|---|---|
| Segmented neutrophil | Vein | 1.3977 | 1 | 0.237114 |
| | **Species** | **6.7332** | **1** | **0.009464**** |
| | Bleeding order | 0.8609 | 1 | 0.353479 |
| | Vein:Species | 1.3771 | 1 | 0.240600 |
| NL ratio | Vein | 0.0858 | 1 | 0.76954 |
| | **Species** | **6.8509** | **1** | **0.00886**** |
| | Bleeding order | 0.5112 | 1 | 0.47464 |
| | Vein:Species | 0.6443 | 1 | 0.42217 |

**Table S8:** Summary of GLMMs that include time between capture and blood collection (holding time), using type II ANOVA tests.

| Cell Type | Variable | $\chi^2$ | df | *p* |
|---|---|---|---|---|
| RBC | Vein | 0.7374 | 1 | 0.3905 |
| | Species | 0.0000 | 1 | 0.9976 |
| | Holding time | 0.0727 | 1 | 0.7875 |
| | Vein:Species | 1.7814 | 1 | 0.1820 |
| WBC | Vein | 1.2387 | 1 | 0.26573 |
| | Species | 1.6937 | 1 | 0.19311 |
| | Holding time | 2.8293 | 1 | 0.09256 |
| | Vein:Species | 0.5079 | 1 | 0.47604 |
| RET | Vein | 3.3149 | 1 | 0.068653 |
| | **Species** | **10.8075** | **1** | **0.001011**** |
| | Holding time | 3.0139 | 1 | 0.082554 |
| | Vein:Species | 0.0075 | 1 | 0.931082 |
| Neutrophils | Vein | 1.4294 | 1 | 0.2319 |
| | **Species** | **16.8495** | **1** | **$4.046 \times 10^{-5}$***** |
| | Holding time | 2.4150 | 1 | 0.1202 |
| | Vein:Species | 2.3059 | 1 | 0.1289 |
| Lymphocytes | Vein | 0.4825 | 1 | 0.4873 |
| | **Species** | **19.6336** | **1** | **$9.381 \times 10^{-6}$***** |
| | Holding time | 0.8977 | 1 | 0.3434 |
| | Vein:Species | 1.6873 | 1 | 0.1940 |
| Monocyte | Vein | 0.1375 | 1 | 0.7108 |
| | Species | 1.3700 | 1 | 0.2418 |
| | Holding time | 0.0195 | 1 | 0.8890 |
| | Vein:Species | 0.3366 | 1 | 0.5618 |

| | | | | |
|---|---|---|---|---|
| Basophil | Vein | 0.4972 | 1 | 0.480716 |
| | **Species** | **3.9032** | **1** | **0.048195*** |
| | **Holding time** | **6.7332** | **1** | **0.009464**** |
| | Vein:Species | 0.0275 | 1 | 0.868367 |
| Eosinophils | Vein | 0.3177 | 1 | 0.57299 |
| | **Species** | **4.4684** | **1** | **0.03453*** |
| | **Holding time** | **6.4316** | **1** | **0.01121*** |
| | Vein:Species | 0.7230 | 1 | 0.39515 |
| Banded neutrophil | Vein | 0.5226 | 1 | 0.4697 |
| | Species | 1.0333 | 1 | 0.3094 |
| | Holding time | 0.6563 | 1 | 0.4179 |
| | Vein:Species | 0.7896 | 1 | 0.3742 |
| Segmented neutrophil | Vein | 0.9708 | 1 | 0.3245 |
| | **Species** | **18.4479** | **1** | **$1.746 \times 10^{-5}$***** |
| | Holding time | 2.4115 | 1 | 0.1204 |
| | Vein:Species | 1.8381 | 1 | 0.1752 |
| NL ratio | Vein | 0.0469 | 1 | 0.8284935 |
| | **Species** | **12.5730** | **1** | **0.0003914***** |
| | Holding time | 0.9022 | 1 | 0.3421922 |
| | Vein:Species | 0.6069 | 1 | 0.4359683 |
